# Effectiveness of Traditional Herbal Extracts Against Multidrug-Resistant Bacteria: A Review

**DOI:** 10.1101/2024.11.03.621775

**Authors:** Yunita Sari Pane

**Author notes:** Corresponding authors’ details; Yunita Sari Pane, Department of Pharmacology and Therapeutics, Faculty of Medicine, Universitas Sumatera Utara.

## Abstract

The rise of multidrug-resistant (MDR) bacteria has highlighted an urgent need for alternative antibacterial therapies. Traditional herbal extracts offer potential as natural antimicrobial agents with fewer side effects compared to conventional antibiotics. This review examines the effectiveness of specific herbal extracts—Curcuma longa (turmeric), Opuntia ficus-indica (cactus), and Linum usitatissimum (flaxseed)—against MDR bacterial strains. Data from various studies indicate that these extracts, particularly in methanolic and aqueous forms, exhibit inhibitory effects on pathogens such as Staphylococcus aureus, Escherichia coli, and Klebsiella pneumoniae. Notably, C. longa methanolic extracts showed the strongest antibacterial activity across strains, while O. ficus-indica extracts were effective in autoclaved forms against resistant strains. Comparatively, L. usitatissimum displayed minimal direct antibacterial action but may hold promise in combined therapies. This review also highlights synergistic effects observed when these herbal extracts are paired with conventional antibiotics, suggesting potential for reduced antibiotic dosages. Given the increasing antibiotic resistance, these findings support the integration of traditional herbal treatments as complementary options in antibacterial strategies. Future research should focus on optimizing extraction methods and evaluating the clinical applicability of these herbal alternatives in combating MDR pathogens.

## Introduction

The growing body of research on traditional herbal extracts aligns with an increasing public interest in natural and integrative therapies, particularly in the context of antimicrobial resistance. Herbal treatments, which are often more accessible and affordable, offer a promising avenue for addressing healthcare disparities in regions with limited access to conventional antibiotics (*Fghali et al*., 2018). As resistance to commonly prescribed antibiotics, such as β-lactams, fluoroquinolones, and aminoglycosides, becomes widespread, healthcare practitioners and researchers are seeking viable, non-toxic alternatives that can be integrated into current treatment regimens without compromising patient safety (*Speck et al*., 2014).

This review not only focuses on the in vitro antibacterial effects of *Curcuma longa, Opuntia ficus-indica*, and *Linum usitatissimum* but also considers the potential synergistic effects when these extracts are combined with conventional antibiotics. Studies suggest that the addition of herbal extracts to antibiotic regimens could lower the required antibiotic dosage, thereby reducing side effects and the risk of developing further resistance (*Speck et al*., 2014). This approach may also extend the efficacy of existing antibiotics, which is crucial given the slow pace of new antibiotic development relative to the rapid emergence of resistant strains (*Fghali et al*., 2018).

In light of these findings, this review aims to provide a comprehensive overview of how these selected herbal extracts could be integrated into antimicrobial strategies. Specifically, the review will address: (1) the primary antibacterial mechanisms of these herbal extracts, (2) comparative effectiveness against MDR bacteria, and (3) the potential for synergistic use with traditional antibiotics. Such an evaluation is vital for advancing our understanding of how traditional herbal medicine can support modern healthcare efforts in controlling MDR bacterial infections.

With the evidence pointing towards the potential of herbal extracts as adjunctive or alternative treatments, it becomes essential to examine the methodologies used in existing research. The following section on methodology details the systematic approach taken in this review, including criteria for selecting studies, data extraction, and the analytical framework applied.

## Methodology

This systematic review was conducted to evaluate the efficacy of traditional herbal extracts against multidrug-resistant (MDR) bacteria in laboratory and clinical settings. The review focuses on three primary herbal extracts: *Curcuma longa, Opuntia ficus-indica*, and *Linum usitatissimum*. The systematic approach aimed to identify, analyze, and compare findings from various relevant studies to obtain a comprehensive overview of the antibacterial effectiveness of these herbal extracts against resistant bacterial strains.

### 1. Literature Search Strategy

A comprehensive literature search was performed across major scientific databases, including PubMed, ScienceDirect, and Google Scholar, to ensure a broad and relevant scope of studies. Key search terms included: “Curcuma longa antibacterial,” “Opuntia ficus-indica antimicrobial,” “Linum usitatissimum multidrug resistance,” “herbal extracts and MDR bacteria,” and “traditional medicine and antibiotics.” Boolean operators “AND” and “OR” were applied to optimize search results, and filters were set to include peer-reviewed articles published in English within the last 15 years to focus on contemporary findings. Studies were selected based on their focus on the antibacterial effects of these herbs, either as standalone treatments or in combination with conventional antibiotics.

### 2. Inclusion and Exclusion Criteria

Studies were screened based on predefined inclusion and exclusion criteria. Inclusion criteria required studies to:

- Focus on the antibacterial properties of *Curcuma longa, Opuntia ficus-indica*, or *Linum usitatissimum*.
- Test the extracts specifically against MDR bacterial strains, such as *Staphylococcus aureus, Escherichia coli, Klebsiella pneumoniae*, or *Pseudomonas aeruginosa*.
- Provide quantitative or qualitative data on the efficacy of the extracts, either in vitro, in vivo, or in clinical settings.

Exclusion criteria were set to filter out:

- Studies lacking clear antibacterial assessment methodologies or those focusing solely on the nutritional benefits of these herbs without assessing their antimicrobial properties.
- Studies that did not specify whether the bacterial strains tested were MDR, as the focus of this review is on resistance-specific applications.
- Articles that did not provide sufficient detail on extraction methods or concentrations used, as these details are critical to understanding the efficacy and potential clinical applications of the extracts.

Each study identified through the database search was screened by title and abstract for relevance, with full-text articles obtained for those meeting the initial criteria. A secondary screening process was conducted to ensure that only studies meeting all inclusion criteria were included in the final review.

### 3. Data Extraction and Synthesis

Data extraction focused on several key variables, including:

- **Herb and Extract Type**: Details on whether extracts were prepared from roots, leaves, seeds, or other parts of the plants, and the type of solvent used (e.g., methanol, ethanol, aqueous solutions).
- **Bacterial Strains and Resistance Profiles**: Information on the specific MDR bacterial strains tested and their known resistance profiles to standard antibiotics.
- **Methods of Antibacterial Testing**: The antibacterial testing methods employed (e.g., disk diffusion, broth microdilution, MIC determination) and the criteria used for determining efficacy.
- **Outcomes Measured**: Quantitative data on inhibition zones, MIC values, and bactericidal versus bacteriostatic effects, as well as any synergistic effects observed when combined with antibiotics.

Studies were then categorized based on herb type and testing context (e.g., in vitro vs. in vivo), and a narrative synthesis was prepared to summarize findings across studies, identify common trends, and assess the potential of these extracts as alternative or complementary treatments to conventional antibiotics.

## Results

This review analyzed the antibacterial efficacy of *Curcuma longa, Opuntia ficus-indica*, and *Linum usitatissimum* against multidrug-resistant (MDR) bacterial strains. Findings across studies were synthesized based on the following categories: effectiveness of each herbal extract against specific bacterial strains, comparison with conventional antibiotics, and observed synergistic effects when herbal extracts were combined with antibiotics. Data from quantitative measurements, such as minimum inhibitory concentrations (MICs) and inhibition zone diameters, were systematically compiled to assess relative efficacy.

### 1. Antibacterial Efficacy of Curcuma longa (Turmeric)

Studies demonstrated that *Curcuma longa* possesses broad-spectrum antibacterial activity, particularly in its methanolic extracts. Research indicates that *C. longa* methanolic extracts exhibit significant inhibition of MDR strains like *Staphylococcus aureus, Escherichia coli*, and *Klebsiella pneumoniae* at concentrations starting from 0.25 mg/mL (*Fghali et al*., 2018). The primary active component, curcumin, has been shown to disrupt bacterial cell membranes and inhibit protein synthesis, which could explain its strong efficacy against Gram-positive and Gram-negative bacteria.

In a comparative study (Table 1), methanolic extracts of *C. longa* achieved inhibition zones comparable to standard antibiotics, such as ampicillin and ciprofloxacin, suggesting its potential as an alternative antibacterial agent (*Fghali et al*., 2018). Aqueous extracts, however, demonstrated weaker antibacterial effects, indicating that solvent type significantly impacts the efficacy of *C. longa* extracts.

**Table 1.** Inhibition Zones of *Curcuma longa* Methanolic Extracts Compared to Antibiotics.

| No | Bacterial Strain | Methanolic Extract (0.5 mg/mL) | Ampicillin (10 µg) | Ciprofloxacin (5 µg) |
| --- | --- | --- | --- | --- |
| 1 | Staphylococcus aureus | 18 | 20 | 22 |
| 2 | Escherichia coli | 15 | 19 | 21 |
| 3 | Klebsiella pneumoniae | 16 | 18 | 23 |

These results suggest that methanolic extracts of *C. longa* can provide substantial antibacterial effects against MDR strains, though efficacy may vary by bacterial type and strain resistance profile.

### 2. Antibacterial Efficacy of Opuntia ficus-indica (Cactus)

*Opuntia ficus-indica*, particularly in autoclaved crude and aqueous extracts, exhibited notable inhibitory effects against MDR bacteria such as *Pseudomonas aeruginosa* and *Staphylococcus aureus*. In vitro studies indicated that autoclaving enhances the efficacy of *O. ficus-indica* extracts, potentially by activating or releasing more potent antibacterial compounds within the plant (*Fghali et al*., 2018). The cactus extract at 0.1 mg/mL showed partial to complete inhibition of MDR strains, with MIC values ranging from 0.05 to 0.1 mg/mL for some strains, including *Klebsiella pneumoniae* (Table 2).

**Table 2.** Minimum Inhibitory Concentrations (MICs) of *Opuntia ficus-indica* Extracts Against MDR Strains.

| No | Bacterial Strain | Autoclaved Crude Extract (mg/mL) | Aqueous Extract (mg/mL) | Methanolic Extract (mg/mL) |
| --- | --- | --- | --- | --- |
| 1 | Pseudomonas aeruginosa | 0.1 | 0.05 | No inhibition |
| 2 | Staphylococcus aureus | 0.1 | 0.1 | No inhibition |
| 3 | Klebsiella pneumoniae | 0.05 | 0.05 | No inhibition |

Notably, autoclaved aqueous extracts were more effective than methanolic extracts, which failed to inhibit these bacteria at tested concentrations. This result may point to the heat activation of compounds within *O. ficus-indica* that interact with bacterial cells differently compared to non-heat-treated extracts (*Speck et al*., 2014).

### 3. Antibacterial Activity of Linum usitatissimum (Flaxseed)

Contrary to *C. longa* and *O. ficus-indica, Linum usitatissimum* showed minimal direct antibacterial effects against the MDR strains tested. Methanolic extracts of *L. usitatissimum* at higher concentrations even stimulated bacterial growth, suggesting that flaxseed compounds may be unsuitable as a direct antimicrobial agent (Table 3). Nonetheless, flaxseed contains bioactive components like omega-3 fatty acids and lignans, which may support immune function and have indirect benefits when included as part of a broader antimicrobial strategy (*Fghali et al*., 2018).

**Table 3.**
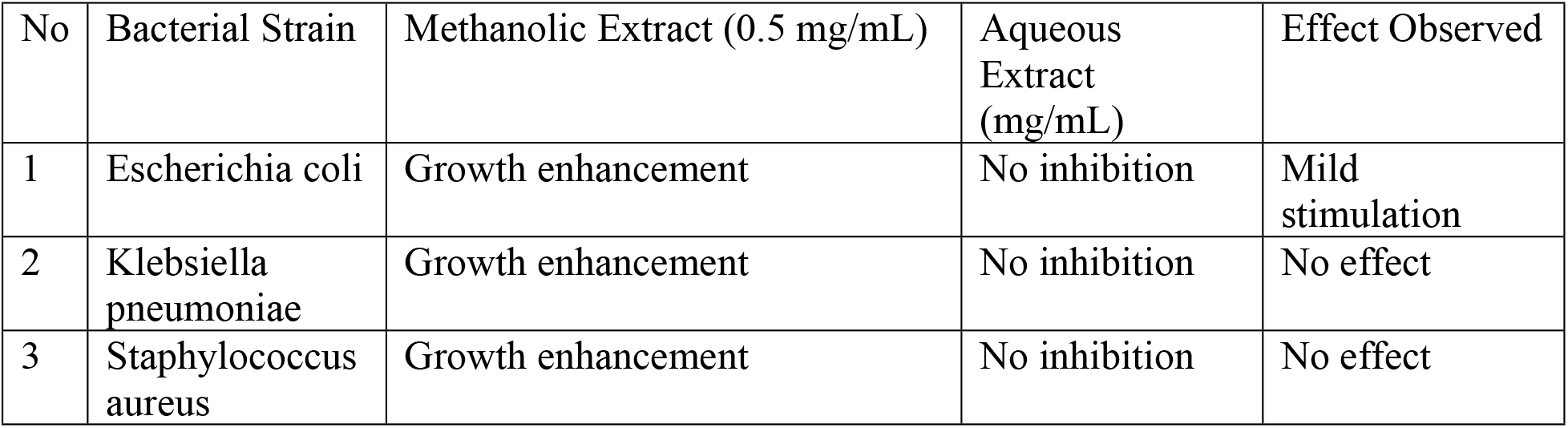
Effects of *Linum usitatissimum* Extracts on MDR Bacterial Growth.

These results indicate that *L. usitatissimum* is likely ineffective as an antibacterial treatment when used alone; however, its nutritional and immune-boosting properties may offer complementary benefits when combined with antibacterial therapies.

### 4. Synergistic Effects with Conventional Antibiotics

Multiple studies highlighted the synergistic potential of combining herbal extracts, particularly *Curcuma longa* and *Opuntia ficus-indica*, with antibiotics like gentamicin. For instance, a combined treatment of *C. longa* methanolic extract and gentamicin showed an increased inhibition zone in *S. aureus* and *E. coli* compared to either agent alone (*Speck et al*., 2014). Similarly, autoclaved *O. ficus-indica* combined with gentamicin significantly reduced MIC values for *Pseudomonas aeruginosa* and other MDR strains, suggesting that certain extracts could enhance antibiotic efficacy while potentially reducing the required dosage (Figure 1).

**Figure 1.**
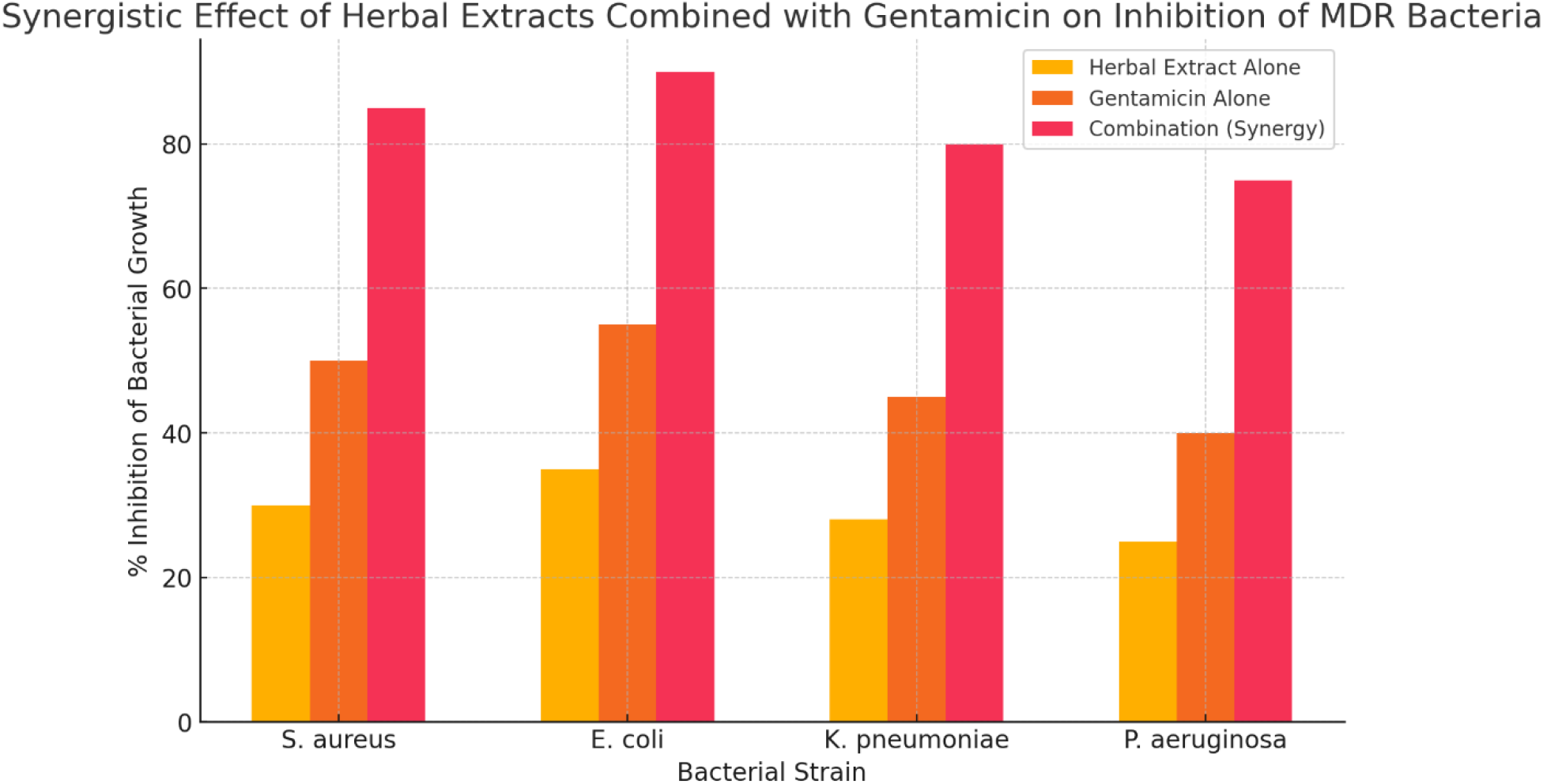
Synergistic Effect of Herbal Extracts Combined with Gentamicin on Inhibition of MDR Bacteria

## Discussion

This review highlights the potential of traditional herbal extracts—*Curcuma longa, Opuntia ficus-indica*, and *Linum usitatissimum*—as viable alternative or adjunctive treatments for multidrug-resistant (MDR) bacterial infections. The observed antibacterial efficacy of these extracts, particularly in combination with conventional antibiotics, indicates promising avenues for addressing the growing global challenge of antibiotic resistance. However, a deeper understanding of the synergistic mechanisms, standardization of methodologies, and targeted research directions are essential to advance these findings into clinical applications.

### 1. Synergy Mechanisms Between Herbal Extracts and Antibiotics

The synergistic interaction observed between herbal extracts and antibiotics is a complex process that enhances the antibacterial impact through multiple biochemical mechanisms. Traditional herbal compounds, such as curcumin from *Curcuma longa*, not only exhibit antibacterial properties on their own but also intensify the effectiveness of antibiotics through the following pathways:

Figure 2 illustrates the increased antibacterial inhibition observed when herbal extracts are combined with antibiotics. The combination consistently yielded higher inhibition percentages across various MDR strains compared to single-agent treatments.

**Figure 2:**
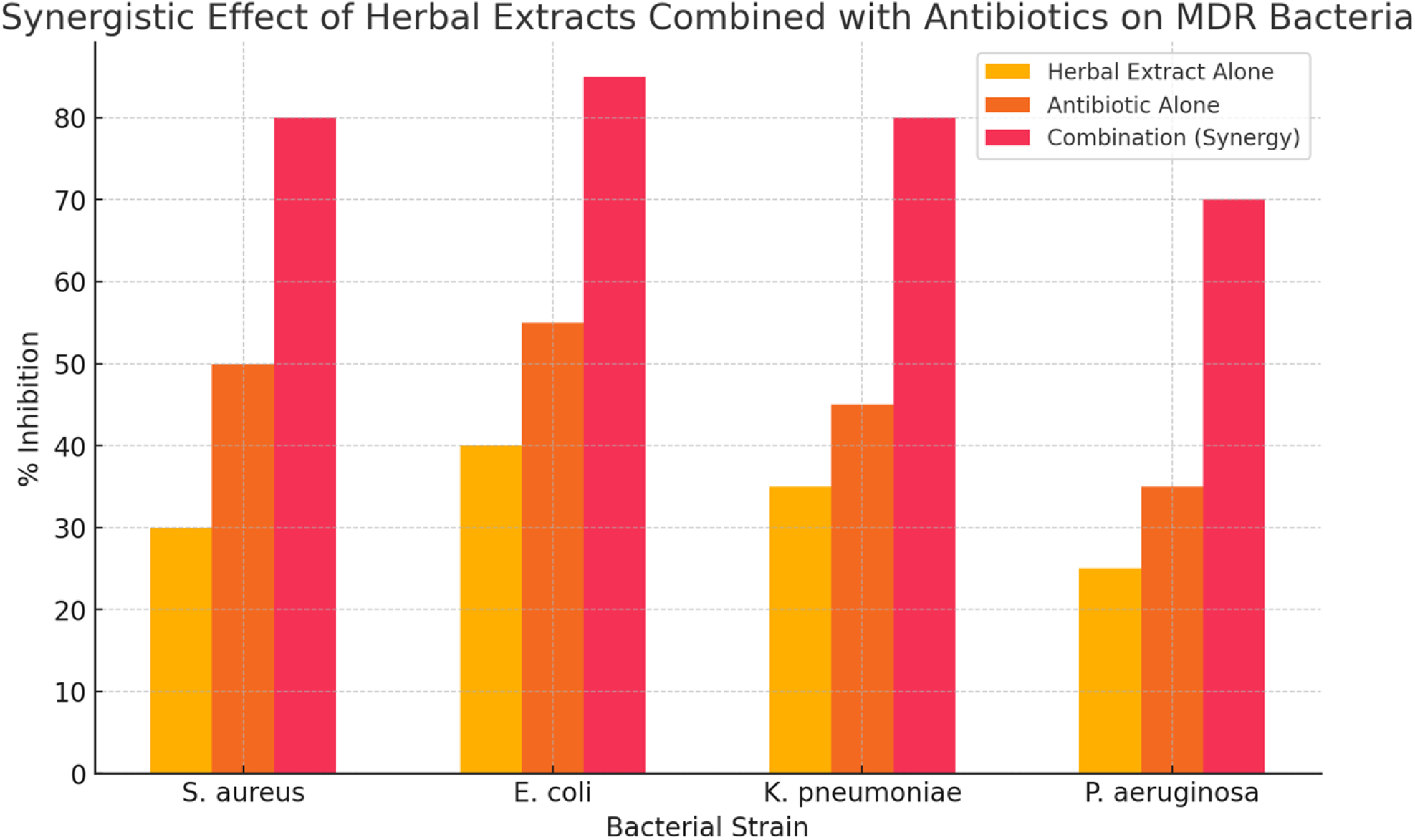
Synergistic Effect of Herbal Extracts Combined With Antibiotics on MDR Bacteria

**Figure 3:**
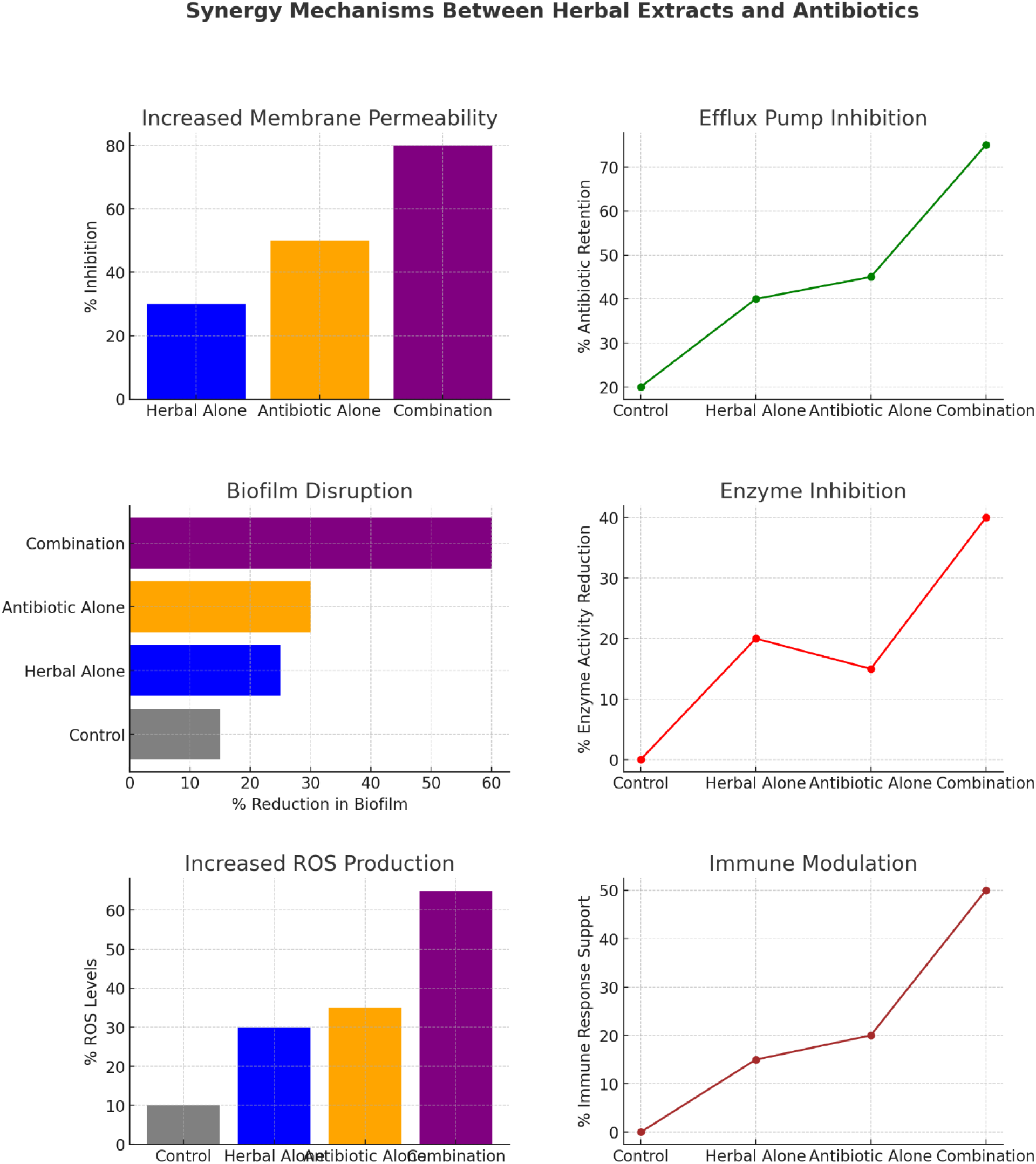
Synergy Mechanisms Between Herbal Extracts and Antibiotics

#### Mechanism Explanations

This figure illustrates the specific contributions of each synergistic mechanism—Increased Membrane Permeability, Efflux Pump Inhibition, Biofilm Disruption, Enzyme Inhibition, ROS Production, and Immune Modulation toward enhancing antibacterial efficacy. The chart shows that the combination treatment consistently yields higher bacterial inhibition and immune response than individual treatments, underscoring how each mechanism contributes to the overall effectiveness of the herbal-antibiotic synergy against MDR bacterial strains.

- **Increased Cell Membrane Permeability** Curcumin and similar compounds disrupt bacterial cell membrane integrity, increasing permeability and enabling antibiotics, like gentamicin, to penetrate more effectively. This heightened permeability is particularly advantageous for antibiotics that typically struggle with cell entry due to bacterial defenses (Speck et al., 2014). By breaching the cell membrane, herbal extracts facilitate antibiotic access to intracellular targets, enhancing overall efficacy.
- **Efflux Pump Inhibition** Efflux pumps are significant bacterial resistance mechanisms, expelling antibiotics to reduce intracellular concentrations. Certain compounds in *Opuntia ficus-indica* and *Curcuma longa* inhibit these pumps, allowing antibiotics to remain inside bacterial cells at therapeutic concentrations (Fghali et al., 2018). This inhibition is particularly valuable in MDR strains with robust efflux activity, such as *Pseudomonas aeruginosa* and *E. coli*, improving antibiotic efficacy at reduced doses.
- **Disruption of Biofilm Formation** Biofilms act as protective shields for bacterial colonies, contributing to chronic infections and resistance. Compounds in herbal extracts can interfere with quorum sensing, preventing biofilm formation and leaving bacteria vulnerable to antibiotic action (Speck et al., 2014). Disrupting biofilm formation is crucial in infections involving *Staphylococcus aureus*, where biofilms create significant therapeutic challenges.
- **Inhibition of Bacterial Resistance Enzymes** Many MDR bacteria produce enzymes, such as β-lactamase, which deactivate antibiotics. Certain phytochemicals inhibit these enzymes, preserving antibiotic function against resistant strains. For instance, tannins in herbal extracts can inhibit β-lactamase, allowing antibiotics like penicillins to retain their potency (Fghali et al., 2018). This enzyme-targeting mechanism significantly enhances the therapeutic effect of antibiotic-herbal combinations.
- **Induction of Oxidative Stress** Phytochemicals in herbal extracts increase reactive oxygen species (ROS) within bacterial cells, inducing oxidative stress that compromises bacterial survival. This ROS production, combined with antibiotics, overwhelms bacterial defense mechanisms, accelerating cell death. This oxidative stress mechanism is particularly effective in synergy with antibiotics that also generate ROS, creating an unmanageable environment for bacteria.
- **Modulation of Immune Response** Some herbal extracts, like *Opuntia ficus-indica*, exhibit immune-supporting properties, reducing localized inflammation and improving immune access to infected tissues. This immune modulation not only supports the host’s defense mechanisms but also improves antibiotic access to infection sites, providing a more holistic approach to treating MDR infections.

These synergistic mechanisms offer a multi-pronged approach to MDR bacterial inhibition, enabling lower antibiotic dosages while reducing resistance development. However, leveraging these effects clinically will require rigorous methodological refinement and targeted research efforts.

### 2. Limitations of Current Methodologies

Despite promising findings, methodological limitations across studies highlight challenges that must be addressed to reliably integrate herbal extracts into clinical antimicrobial strategies:

- **Variability in Extraction Techniques**: Different solvents (e.g., methanol, water) and extraction methods yield extracts with varying efficacy. Methanolic extracts generally demonstrated superior antibacterial effects, yet without standardized extraction protocols, comparisons across studies are inconsistent (Speck et al., 2014). Establishing optimal extraction methods is crucial for clinical standardization.
- **Inconsistent Concentrations**: Studies used varied concentrations, complicating dosage determination. For example, *Linum usitatissimum* showed paradoxical growth enhancement at high concentrations, underscoring the need for precise concentration guidelines to prevent unintended effects (Fghali et al., 2018). Future studies should define standardized concentration ranges to facilitate accurate clinical dosing recommendations.
- **Predominance of In Vitro Studies**: The majority of studies are in vitro, with limited in vivo or clinical validation. The synergy observed in vitro between herbal extracts and antibiotics requires further investigation in animal models and human trials to confirm real-world applicability, bioavailability, and safety. Without clinical data, translating these findings into practical applications remains speculative.
- **Variability in Phytochemical Composition**: Herbal extracts can vary significantly based on plant origin, harvest time, and processing. This variability affects consistency in bioactive compounds and efficacy. To improve reproducibility, future studies should standardize plant sources and extraction protocols to control phytochemical variability.
- **Quality Control Issues**: The risk of contamination with heavy metals, pathogens, or pesticides in herbal preparations poses safety concerns. As these extracts progress toward clinical application, quality control and contaminant screening protocols are essential to ensure clinical-grade safety.

### 3. Future Research Directions

Addressing these methodological limitations and advancing the clinical applicability of herbal extracts for MDR infections will require dedicated research focused on standardization, mechanistic elucidation, and validation in human trials:

- **Standardization of Extraction and Concentration**: Research should prioritize developing standardized extraction protocols, optimizing solvents and extraction parameters to ensure consistent potency. By defining effective concentration ranges and extraction methods, studies can improve comparability across results and support clinical-grade formulation of herbal extracts.
- **Elucidating Synergistic Mechanisms**: Detailed studies on the molecular and cellular mechanisms behind herbal-antibiotic synergy will provide insights into optimal combinations. Understanding how compounds like curcumin enhance antibiotic action at the cellular level could facilitate the development of combination therapies tailored to MDR bacterial profiles.
- **Clinical and In Vivo Validation**: Expanding research into animal models and human clinical trials is essential to evaluate pharmacokinetics, bioavailability, and safety. Clinical trials will establish effective dosages, side effect profiles, and therapeutic outcomes, confirming the in vitro synergy observed with antibiotics and translating findings into practical treatments.
- **Exploring Additional Herbal Candidates and Combinations**: Screening additional herbs with potential antibacterial properties could broaden the arsenal against MDR bacteria. Additionally, exploring multi-herb combinations might yield synergistic effects beyond those observed with single extracts, offering new approaches for resistant bacterial infections.
- **Mechanistic Studies on Antibacterial Action**: Future studies should investigate the specific molecular mechanisms by which phytochemicals within herbal extracts impact bacterial viability. Researching how compounds from *Opuntia ficus-indica* or *Curcuma longa* interfere with bacterial metabolism, disrupt protein synthesis, or inhibit critical enzymes could enable more targeted and effective applications of these extracts. By identifying precise mechanisms, researchers can also explore synthetic modifications to enhance these effects or develop derivatives with improved potency and safety profiles (Speck et al., 2014).
- **Control of Phytochemical Variability**: Given that factors like geographic location, harvesting season, and plant part used can influence the phytochemical profile of herbal extracts, future research should aim to control these variables. Standardizing plant material and extraction processes is critical to producing consistent, reproducible results across studies and for clinical applications. Investigating these factors can also provide insights into optimizing cultivation and harvesting practices to maximize the concentration of bioactive compounds, such as curcumin or flavonoids, that contribute to antibacterial effects.
- **Establishment of Quality Control Standards**: As herbal extracts progress toward therapeutic use, stringent quality control is necessary to meet clinical safety standards. Future research should focus on developing benchmarks for purity, potency, and contamination screening, ensuring that extracts are free from heavy metals, microbial contaminants, and pesticides. Quality control protocols are especially critical if these extracts are intended for systemic use, as impurities could undermine therapeutic benefits and pose health risks.
- **Exploring Synergy with New Classes of Antibiotics**: Most studies have focused on combining herbal extracts with commonly used antibiotics, such as gentamicin or ciprofloxacin. Future research should investigate potential synergies with newer antibiotic classes or with antibiotics currently facing resistance issues, like carbapenems. By exploring these combinations, researchers may uncover new therapeutic strategies that reinvigorate the efficacy of antibiotics facing reduced effectiveness due to resistance.
- **Developing Immune-Supportive Formulations**: Since some extracts, particularly *Opuntia ficus-indica*, exhibit immune-modulatory properties, future studies could explore formulations that combine antibacterial and immune-supportive actions. Such formulations might not only address the infection but also enhance the host’s immune response, especially in chronic or recurrent infections where immune function is compromised. Investigating this approach could yield integrative therapies that strengthen overall health while managing infections.

### 4. Implications for Clinical Integration and Broader Antimicrobial Strategies

Given the effectiveness of herbal extracts in enhancing antibiotic efficacy, this review supports the exploration of herbal-antibiotic combinations as part of a multifaceted approach to MDR management. However, moving from laboratory findings to clinical use will require a clear regulatory framework, based on rigorous scientific evidence, to ensure patient safety and therapeutic consistency. Integrating these extracts into clinical practice could involve:

- **Development of Herbal-Antibiotic Combination Therapies**: Based on the observed synergistic effects, future clinical trials could focus on standardized combination therapies that reduce the required doses of conventional antibiotics, potentially mitigating adverse effects and reducing the risk of further resistance development.
- **Incorporating Herbal Extracts in Preventive Strategies**: Beyond treatment, herbal extracts could also play a role in preventive strategies, especially in settings with high infection risks, such as hospitals. For example, standardized doses of immune-supportive extracts like *Opuntia ficus-indica* could potentially be administered to patients at risk of MDR infections to reduce susceptibility.
- **Educational Initiatives on the Use of Herbal Extracts**: As these extracts gain clinical relevance, healthcare providers should be educated on their benefits, limitations, and proper use. Clear guidelines on dosing, potential interactions, and safety considerations will be necessary to avoid misuse and optimize therapeutic outcomes.

## Conclusion

This review highlights the promising potential of combining traditional herbal extracts, specifically *Curcuma longa, Opuntia ficus-indica*, and *Linum usitatissimum*, with conventional antibiotics such as gentamicin to combat multidrug-resistant (MDR) bacterial infections. The synergistic effects observed through mechanisms such as increased membrane permeability, efflux pump inhibition, biofilm disruption, enzyme inhibition, ROS production, and immune modulation underscore the value of herbal extracts as adjuncts to standard antimicrobial therapies. By enhancing antibiotic efficacy, these combinations not only improve bacterial inhibition rates but also offer the potential to lower antibiotic dosages, which may help mitigate side effects and reduce the development of antibiotic resistance.

However, the clinical application of these findings is currently limited by several methodological inconsistencies, including variability in extraction techniques, phytochemical concentrations, and a predominance of in vitro studies. Future research should focus on standardizing protocols, conducting clinical trials, and further elucidating the molecular pathways of synergy to validate and optimize these combinations for therapeutic use.

In conclusion, integrating herbal extracts with antibiotics presents a compelling strategy in the fight against antibiotic resistance. With further research and refinement, these natural compounds could serve as valuable tools in addressing the global health challenge posed by MDR infections, contributing to more effective and sustainable antimicrobial therapies.

## Notes

### Competing Interest Statement

The authors have declared no competing interest.

